# SOCS1/SOCS3 Immune Axis Modulates Synthetic Perturbations in IL6 Biological Circuit for Dynamical Cellular Response

**DOI:** 10.1101/2020.08.31.276634

**Authors:** Bhavnita Soni, Shailza Singh

**Author notes:** Corresponding author, /95.

## Abstract

Macrophage phenotype plays a crucial role in the pathogenesis of Leishmanial infection. Pro-inflammatory cytokines are the key regulators that eliminate the infection induced by Janus kinase/signal transducer and activator of transcription (JAK/STAT) pathway. Suppressor of cytokine signaling (SOCS) is a well-known negative feedback regulator of JAK/STAT pathway. However, change in expression levels of SOCS in correlation with the establishment of infection is not well understood. Mathematical modeling of IL6 signaling pathway have helped identified the role of SOCS1 in establishment of infection. Furthermore, the ratio of SOCS1 and SOCS3 has been quantified both *in silico* as well as *in vitro*, indicating an immune axis which governs the macrophage phenotype during *L. major* infection. The ability of SOCS1 protein to inhibit the JAK/STAT1 signaling pathway and thereby decreasing pro-inflammatory cytokine expression makes it a strong candidate for therapeutic intervention. Using synthetic biology approaches, peptide based immuno-regulatory circuit have been designed to target the activity of SOCS1 which can restore pro-inflammatory cytokine expression during infection.

## INTRODUCTION

Suppressor of cytokine signaling (SOCS) is known for negative feedback regulation of Janus kinase/signal transducer and activator of transcription (JAK/STAT) pathway(1). In various infectious disease, SOCS regulates the activation of cell by pro-inflammatory cytokines (2). Post natal death of SOCS1 knockdown mice have been reported within three weeks due to IFN-γ-induced hyper-inflammation (3). Contrary to this, there are few infectious diseases wherein parasite inhibits activation of immune cells through selective expression of few SOCS (4). One such example is Leishmaniasis wherein expression of SOCS isoforms plays a crucial role in the establishment of infection (5).

We all know the immune system is a tremendously complex system which needs to be understood through mathematical models of varied and myriad number of interconnecting components. Intuitively, systems biologists talk in two scenarios wherein one focuses on intracellular molecular networks involved in gene regulation, signaling and the other molecular processes while the other focuses on systemic aspects of immune system dynamics. Using systems biology approaches, we have already established mathematically, the role of SOCS1/SOCS3 ratio in raising the immune response during early stage of *L. major* infection (6) (BIOMD0000000873). We observed the ratio to be >1, which depicts that an incremental increase in concentration of SOCS1 eventually shuts down the pathways responsible for its pro inflammatory behavior. One of the key strength of this approach is that it allows simultaneous validation of the observations obtained from mathematical modeling and mimics *in vitro/in vivo* systems. To commensurate this, modeling and simulations are integral part of systems biology, where in, mathematical modeling guides’ *in vitro/ in vivo* experimentation which further aid in model refinement leading to better understanding of complex biological systems. Thus, model refinement is an important step toward unfolding the crucial dynamics of complex biological systems (7). It would be worthwhile to mention here that performing *in silico* deterministic simulation has much more advantage than the stochastic one. In deterministic model, a given choice of parameters and initial conditions always lead to the same set of model predictions, models of this sort typically are in the form of coupled ODEs describing the dynamics of molecular concentrations as most appropriate. However, investigation of a stochastic model is complicated by the fact that system trajectories varies from realization to realization and solving the master equation becomes little more complex due to the combinatorial explosion of the parameter and configuration space(8). In corroboration to this, based on literature evidences we have constructed comprehensive signaling network of cytokine IL-6. We know IL6 involvement in a multitude of processes right from immune response to pathogens to cancer related processes and also in the regulation of inflammatory processes linked to insulin resistance(9, 10, 11). Analysis of the network suggests several model modifications in order to better fit available knowledge and data, which further helped intrigued our experimental hypothesis to be pursued. Thus, in the present work, using experimental observations we have refined our previous models of Healthy state model (HSM) and Diseased state model (DSM) established during early stage of *Leishmania major* infection. HSM referring to M1 and DSM refers to M2 phenotype of macrophage respectively during infection.

Later the refined models were used to design therapeutics based on synthetic biology approaches using biological components (promoter, RBS, RNA polymerase, spacer etc.) that have been rewired to act as a transcriptional pool. Each biological entity represents one transcription unit. Extensive work has been done using tunable synthetic circuit in mammalian cell for therapeutic purpose such as the use of gene circuit for sensing and suppressing inflammation (12), treatment of metabolic syndrome (13), for anti- cancerous gene therapy (14) and the use of synthetic gene circuit for immune mediated therapy (15). Besides these, synthetic mammalian gene circuit has also been used to deliver specific RNA to cell (16). The success of the mammalian based synthetic gene regulatory circuit in various diseases has drawn our attention and motivated us to generate a potent novel therapeutics in Leishmaniasis. Here, in this paper, we have assimilated different parts of the transcriptional unit (promoter, RBS, RNA polymerase, spacer etc.), pooled from genetic pool to generate a functionally active synthetic circuit. The design is precise keeping it modular in fashion which ensures the simple and reproducible workability of the circuit for lest it should not be visualised as Rube Goldberg machine. To the best of our knowledge, this manuscript serves as the first ever report of IL6 based synthetic gene regulatory circuit for treating *L. major* infection at cellular level.

There are various forms of synthetic devices used for therapeutic purpose, more complex types are oscillators and toggle switches which contains two or more stable states with or without an intermediate unstable states (17). The simpler form of synthetic device is the repressilator, characterized by the presence of feedback loop with at least three genes out of which one encode for protein that represses the next gene in loop (18). In the present study, we have used the combination of toggle switch and repressilator to design the synthetic circuit that may tweak the immune response from Th2 to Th1 type during early stage of leishmanial infection. The immuno based synthetic device serves as the first attempt to revert the anti-inflammatory action of IL6 into its pro-inflammatory behavior through mathematically established SOCS1/SOCS3 immune axis. Thus, the increasing body of vast knowledge together with comprehensive mathematical analysis may aid immuno-based synthetic device to become a reality in Leishmania infection model system (Fig.1).

**Fig.1.**
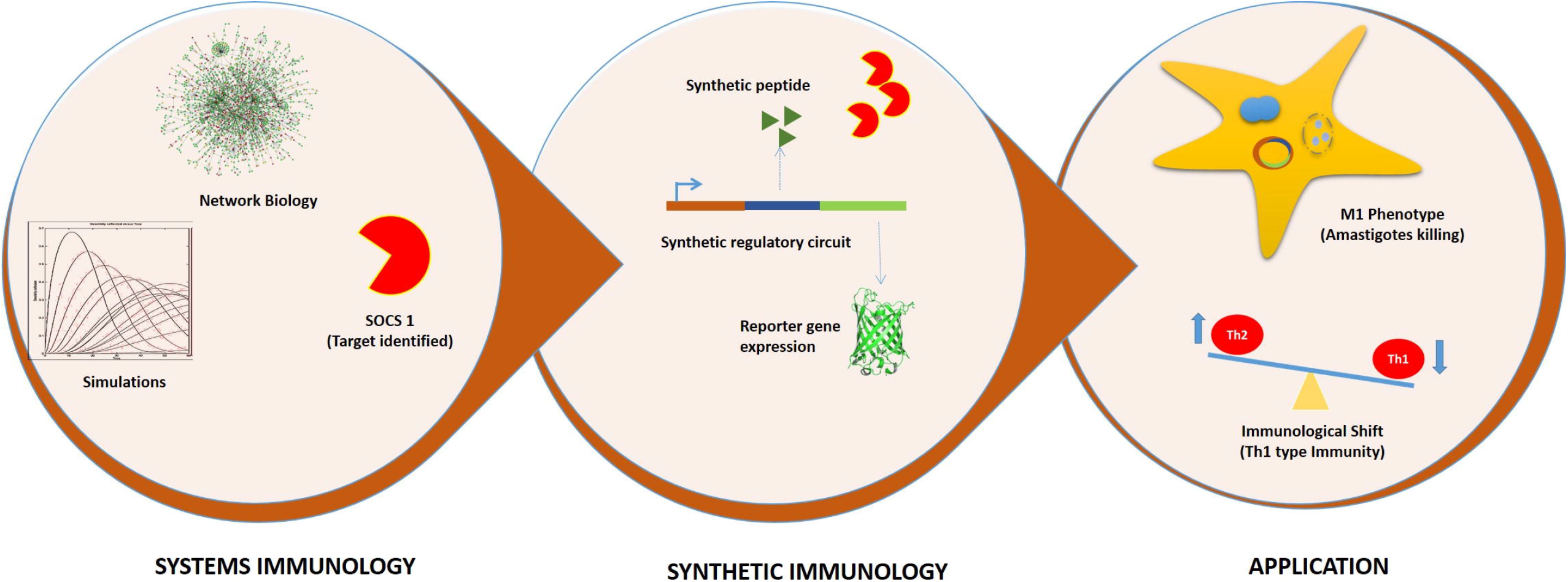
Graphical representation of Systems and Synthetic immunology

## RESULTS

### A. SOCS1/SOCS3 differential expression governing macrophage polarization

Interleukin 6 cytokine is one of the major cytokine which is released during interaction of LPG (Lipophosphoglycan) with TLR2 at an early stage of infection (19). Previously, we have generated two models deciphering the dual action of IL6 in Leishmaniasis i.e. DSM depicting the anti-inflammatory role and HSM showing pro inflammatory role of IL6 (6) We have hypothesized that IL6 may act as an anti-inflammatory cytokine causing selective expression of SOCS proteins resulting in converting the macrophage in M2 phenotype during Leishmanial infection. In this process, first the amount of IL6 is quantified during post one hour of *L. major* infection (Fig.2g) and as per our previous observation, 100 time unit simulation of the diseased state model shows differential expression of SOCS1 and SOCS3 protein (6). This crucial finding was further validated in *in vitro* by infecting the macrophages with *L. major* stationary phase promastigotes in the presence of Interleukin 6 cytokine. Western blot data depicted increased expression of SOCS1 and SOCS3 protein during infection which further got enhanced in presence of IL6 treatment. Densitometric analysis identifies the ratio of SOCS1/SOCS3 as 3:2 post one hour of infection (Fig.2e and 2f). The obtained data signifies the anti-inflammatory role of IL6 in establishing SOCS1/SOCS3 immune axis during early stage of infection.

**Fig. 2.**
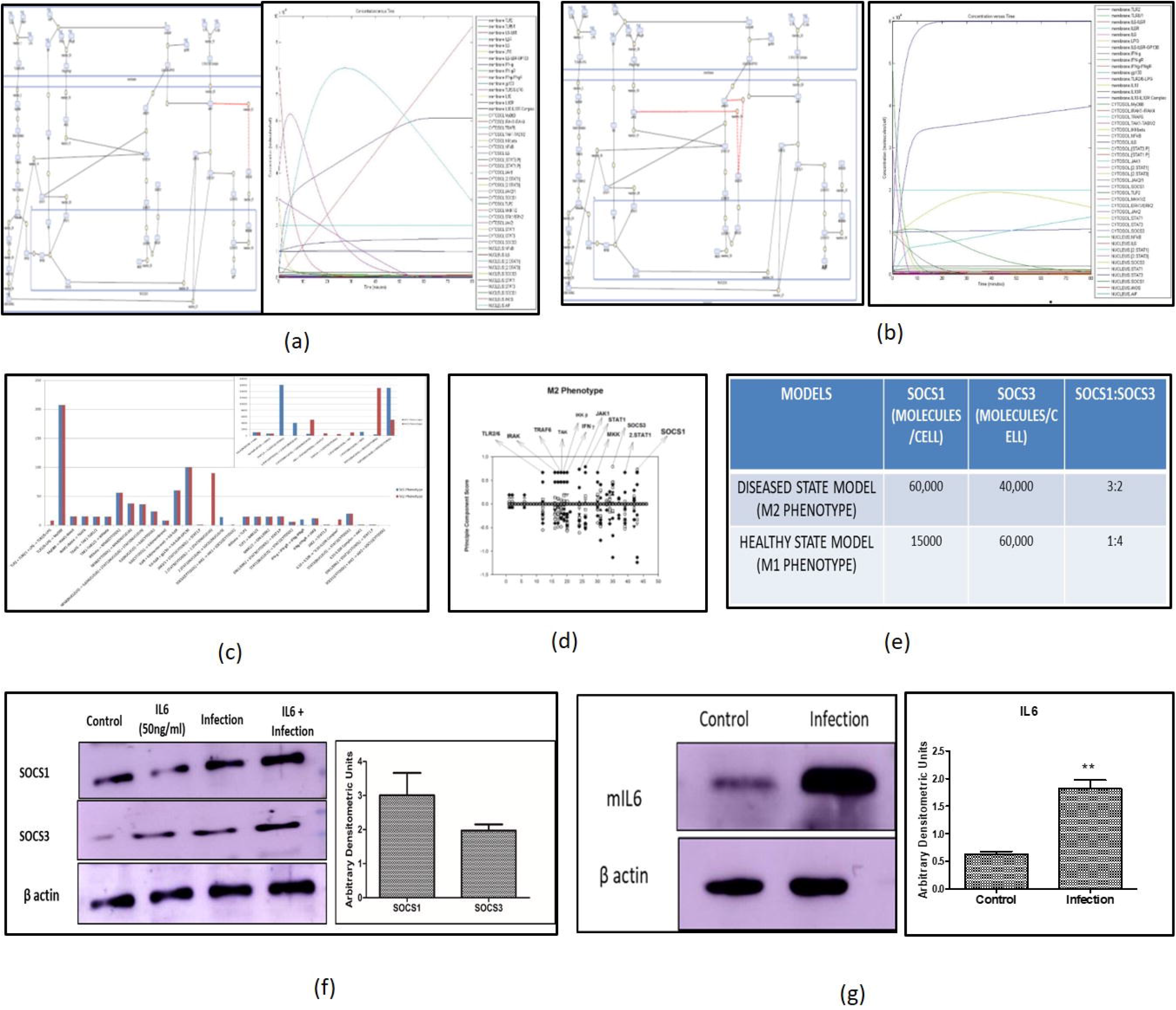
Refinement of mathematical model: (a) Mathematical model of M1 phenotype depicting higher production of iNOS and SOCS3, (b) Mathematical model of M2 phenotype representing higher production of Anti-inflammatory factors (AIF) and SOCS1 at the end of 80 time unit simulation. (c) Comparative flux analysis representing reactions with higher flux. (d) Principle component analysis of DSM model showing key components governing the system. (e) Quantification of SOCS1/SOCS3 concentration ratio obtained after simulation of mathematical model. (f) Further, macrophages were exposed to stationary phase promastigotes in ratio of 1: 10 (macrophage: parasite) for 24 hrs and un-internalized parasite was washed followed by treatment of IL6 (50ng/ml) for 1 hour post infection. Ratio was further quantified through western blotting followed by densitometric analysis of the blots against b actin level. (g) Quantification of IL6 cytokine has been done post one hour of *L. major* infection through western blotting. For all western blot experiments, equal amount of protein was resolved and blotted for b-actin to ensure equal loading. The experiments were performed thrice. Results from one representative experiment are shown. The densitometric value represents mean ± SD with p value *p < 0.05, **p < 0.01, ***p < 0.001.

The IL6 mathematical model is constructed for it to be testable through further experimentation. We need to quantify the uncertainty of those predictions, given the information they are built upon. Either the local analysis of sensitivities or the non-local sampling of parameter space can be used to estimate prediction uncertainties. Model re-parameterizations and targeted experiments can result in identifiable parameters. Here, in this paper what we have seen is that even though some parameter in the ensemble vary considerably, the ensemble of trajectories shows much less variation. Nonetheless, the reality is that many mathematical models are published with parameters that do not systematically fit to the data. We would like to reiterate that our previously published IL6 model (BIOMD0000000873) does not fit the experimental data leading us to only use the synthetic data generated by the model itself. Counter intuitively, we went ahead in understanding the sensitivity of model predictions to parameters which would suggest possible perturbations of interest and adopted the established protocol of our lab.

On the basis of aforementioned strategies, refinement of the mathematical models i.e. Diseased state (M2 phenotype); DSM and Healthy state (M1 phenotype); HSM have been performed. Each model contained 46 species comprising of 41 reactions in DSM and 40 reactions in HSM respectively (S1). After a simulation for 100 time unit, DSM shows increased concentration of Anti-inflammatory factors (AIF) as well as SOCS1 protein whereas HSM shows higher concentration of iNOS and SOCS3 (Fig. 2a and 2b). The model has been submitted to BioModel database with ID MODEL2005140001. Later, both the models were subjected to Sensitivity and Principal component analysis followed by flux analysis. In this case, the PCA approach was laid down to identify multivariate relationship between IL6 signaling events and also link them to *in vitro* cellular phenotypes. We adopted the gold standard method of PCA and mathematical modeling approaches to more accurately differentiate between disease progressions states (Fig 2d). These multivariant analyses led us to churn out the major reactions that adds to disease progression at cellular level (Table1), followed by the Ordinary differential equation of (ODEs) of major biochemical reaction.

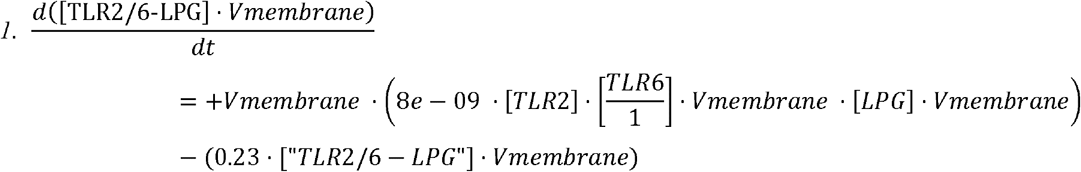

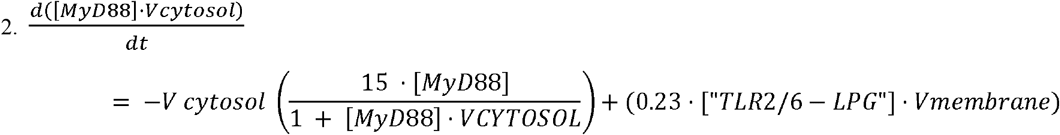

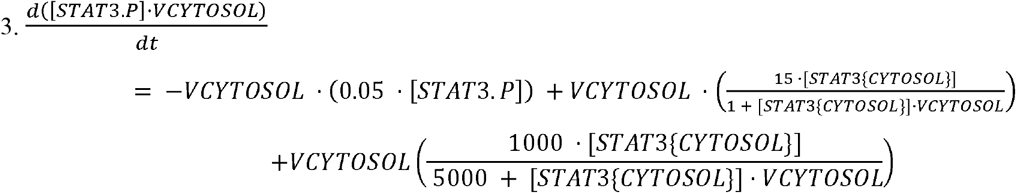

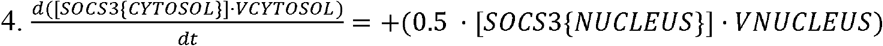

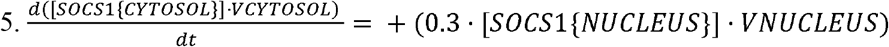

**Table 1.** Reaction that governs the disease progression at cellular level.

### B. Systems study reveals phosphorylated STAT1 and STAT3 as cross talk

At the intracellular level, model captures a variety of signaling events, most important being the signal transduction networks emanating from receptors and engaging downstream in crosstalk. With respect to our previous observation (6)and now in refined models as well, cytoplasmic phosphorylated STAT1 and STAT3 are found to be as cross talk points between TLR2/TLR6-IL6 signaling pathway, later validated through western blotting. We observed that, in both the cases, the activation of either of the two pathways represents phosphorylation of STAT1 and STAT3, and is inhibited with the addition of respective inhibitor (Fig 3c, 3d). Each pathway is then further activated in presence of their respective inhibitors signifying constant phosphorylation of both the STATs (Fig 3b). This shows phosphorylated STAT1 and STAT3 acts as a cross talk point between IL6 and TLR2 signaling pathway (S7).

**Fig. 3.**
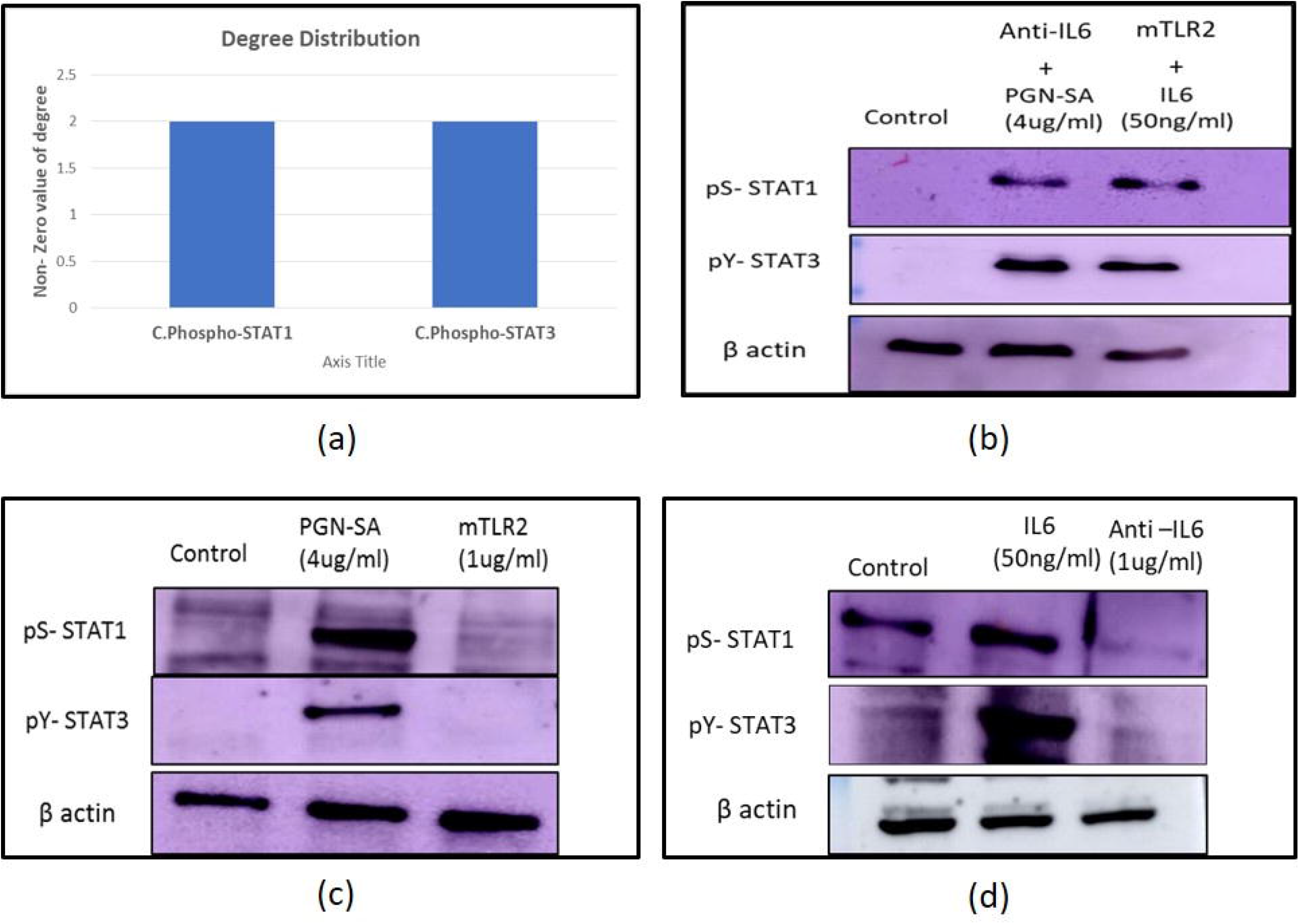
Identification of cross talks between TLR2/TLR6-IL6 signaling pathways (a) Degree distribution of cross talks identified through network analysis (b) Western blotting was used to check expression levels of phosphorylated STAT1 (S727) and STAT3 (Y705) during activation of one pathway & inhibition of another pathway and vice versa. (c) & (d) Expression levels of phosphorylated STAT1 (S727) and STAT3 (Y705) during activation and inhibition of their respective pathway. For all western blot experiment, equal amount of protein was resolved and blotted for b-actin to ensure the equal loading. The experiments were performed thrice. Results from one representative experiment are shown.

### C. Multi objective optimization of mathematical model

The idea of performing multi-objective optimization of mathematical model is to elucidate how network is evolvable with respect to changing environmental condition. The evolved network could be a better platform to generate any kind of therapeutics. *Leishmania* interferes with the IL6 signaling network by modulation of SOCS1:SOCS3 ratio. SOCS1 is responsible for anti-inflammatory behavior and SOCS3 corresponds to pro inflammatory behavior. Once the rewiring of the network is completed, the network should evolve towards pro inflammatory phenotype. Since obtaining evolvability of a system is a multi-optimization problem, we opted for Multi-objective genetic algorithm (MOGA) and defined objective function as :

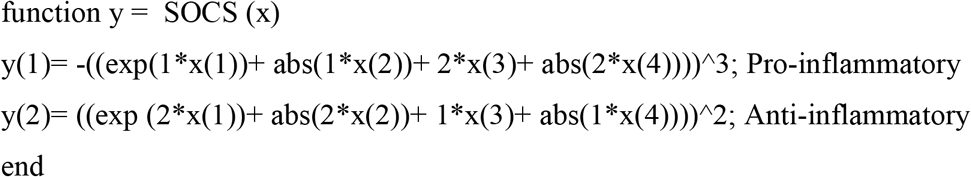

The decision variables embedded in y (1) and y (2) objective functions are cytokines, whose expression levels would be measured in *in vitro* experiments. They are denoted as x1, x2, x3 and x4, representing the cytokines IL-10, TGF β, TNF-α and IFN-γ respectively. It shows that x1 and x2 decision variables represent the anti-inflammatory cytokines for the fitness function of SOCS1 protein (y (1)). Similarly, x3 and x4 represents (pro-inflammatory) fitness function of SOCS3 protein. When genetic algorithm is performed with defined objective functions, a graph of individual v/s generation is obtained showing elite population (Fig 4a), this signifies the importance of SOCS1: SOCS3 ratio, as a character of elite population. The average distance between the individual is low throughout the run, indicate decreased mutation rate or conservedness of the ratio throughout many generations (Fig 4b). The Pareto front obtained for the opposing objective functions have more than 30 non-dominated solutions that are not discontinuous and the average spread measure for these solutions is 0.167776 (Fig 4c &4d). The Pareto optimality obtained using genetic algorithm states that during the process of natural selection the ratio of SOCS1: SOCS3 obtained through mathematical modeling analysis (Fig.2) together with the anti and pro inflammatory cytokine is selected (conserved) and passed onto next generation as an elite character. By targeting this conserved ratio, the designed therapeutics would be more effective for generations turning the macrophage polarization towards M1 phenotype.

**Fig.4.**
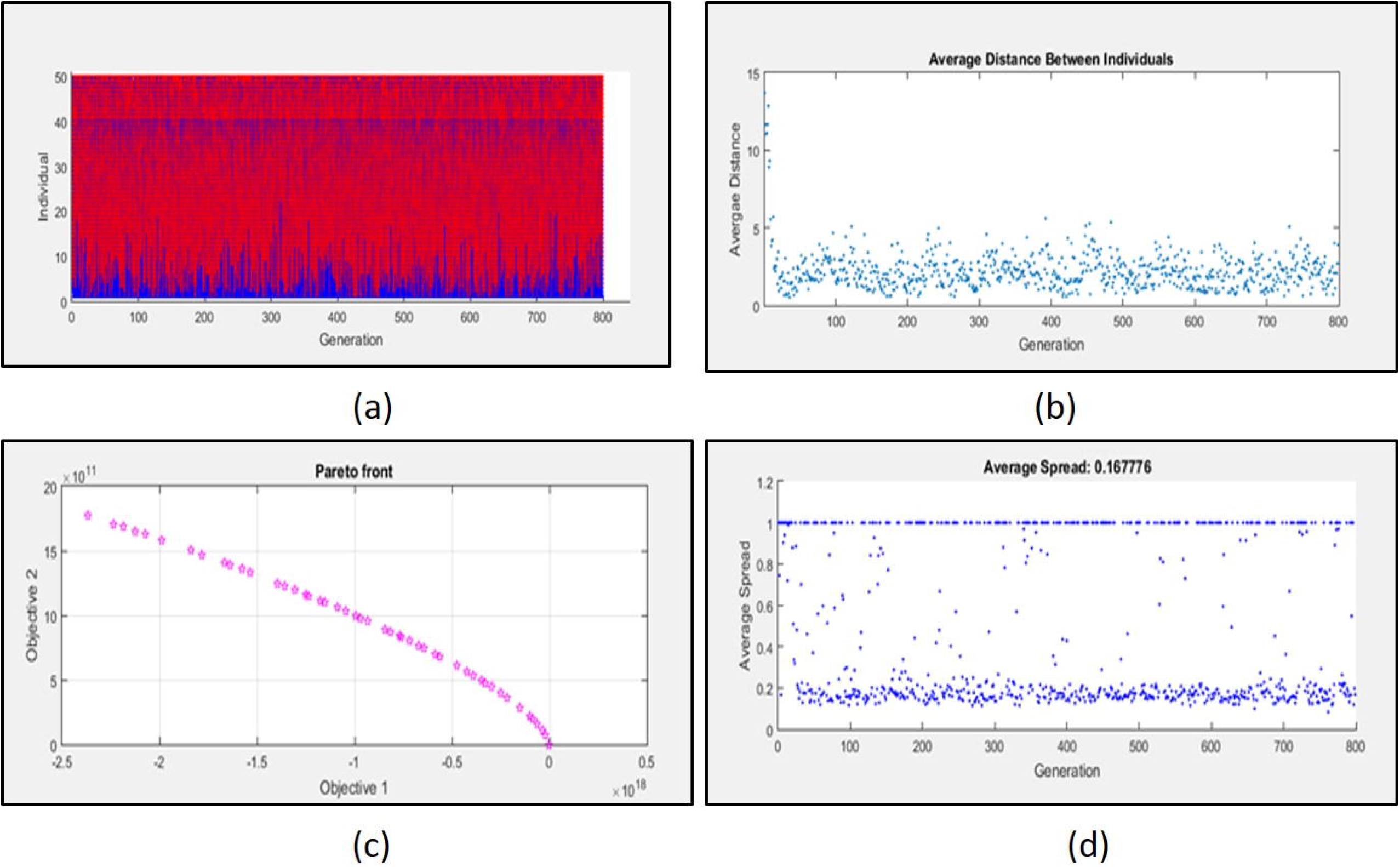
Multi objective genetic algorithm for optimization of SOCS1 /SOCS3 ratio (a) Graph of elite population (b) Graph showing average distance between individuals indicating the decreased mutation rate (c) Pareto front between two objective function representing non dominated solution (d) Graph depicting the average spread measure of the solution.

Evolutionary algorithms iteratively improve circuit performance by randomly mutating parameters across a population of circuits, and retaining circuits with the highest fitness and selectivity. This approach generates functioning circuits with fewer computations and has an added advantage of providing some insight into how gene circuits might evolve during natural selection. The final output of our algorithm is a simplified gene network that defines a dynamical system whose response is a switch-like function of its inputs with the anti-inflammatory response getting shifted towards pro-inflammatory response.

### D. SOCS1 as a target for therapeutic intervention

By performing the process of model reduction, there are various other reactions which have been filtered out in both the models (Table1). One of the major reaction with high flux during analysis was the formation of active cytoplasmic SOCS1 protein (SOCS1 {NUCLEUS} -> SOCS1 {CYTOSOL}) in DSM. Cytoplasmic SOCS1 was among the major nodes identified through principle component analysis and also showed high sensitivity score (Fig 1). Thus, cytoplasmic SOCS1 is selected as a target for further therapeutic intervention.

### E. Peptide Design, Docking and MD simulation of Selected Complex

We designed a small peptide library of 15 peptides based on Machine Learning **(S2A)**, assumption was on the non-conserved region in SOCS1 protein. Peptide 8 (NSQKADDLVDNNVI) was selected on the basis of number of interacting residues as well as low energy complex forming ability **(S2B)**. The SOCS1-Peptide8 complex was then subjected to 30ns MD simulation. The RMSD plot shows that the complex got stabilized post 20 ns and the complex achieved its minimum energy state conformation (Fig.5f). **(S3)**.

**Fig.5.**
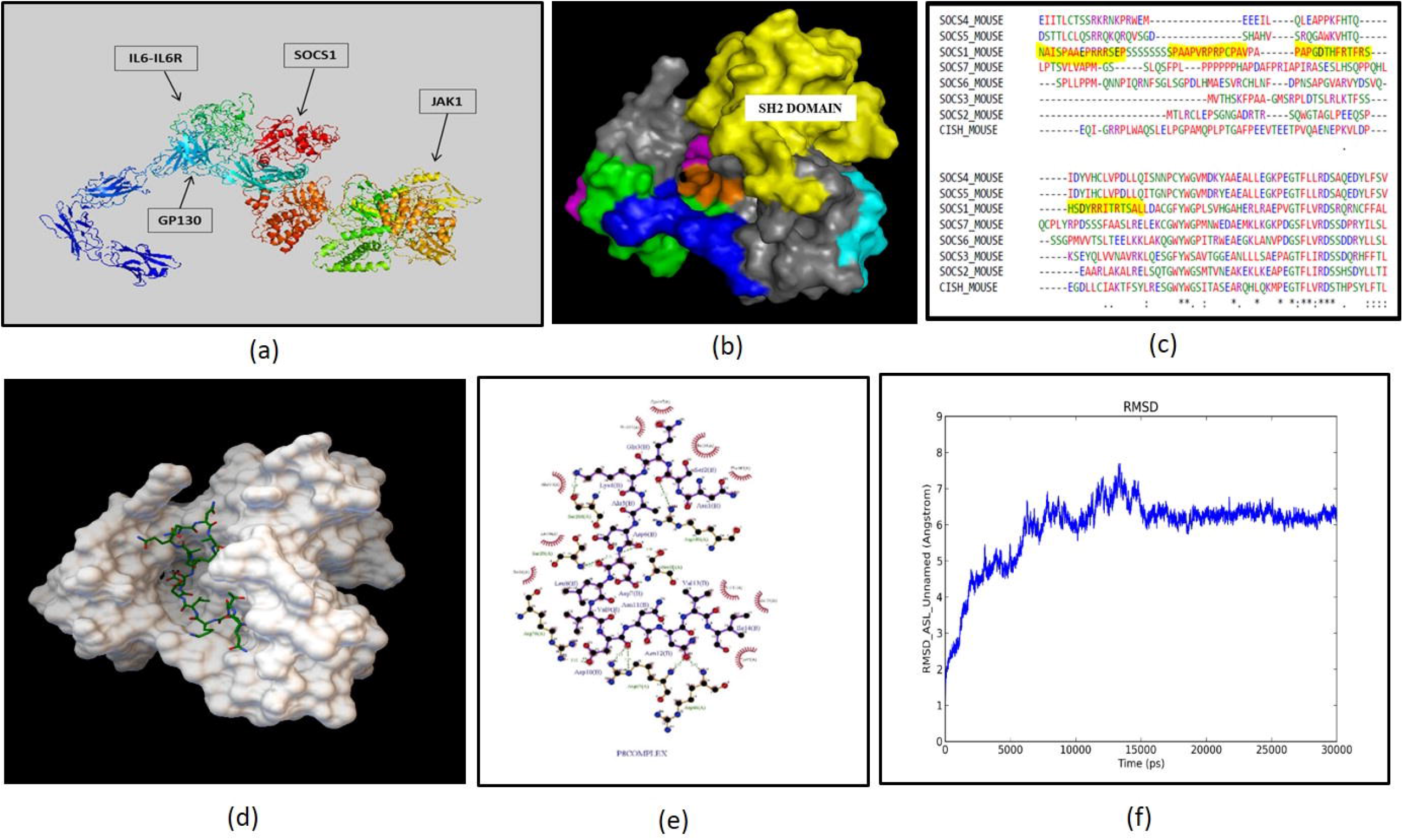
Peptide designed against SOCS1protein: (a) Protein-protein docking of IL6-IL6R-gp130 complex with JAK1 and SOCS1 to identify interfacial residue during interaction. (BioLuminate Package, Schrödinger Release 2017-3 Suites). (b &c) Multiple sequence alignment identified non-conserved regions near SH2 domain of SOCS1 protein as a target for peptide design. (d) The complex shows that peptide 8 has been docked at desired region near SH2 domain **t**hat may inhibit the interaction of SOCS1 and gp130 (Autodock v1.5.6) (e) Interaction plot of SOCS1 and Peptide8 amino acid residues (Ligplot+). (C) Molecular dynamics simulation of SOCS1-peptide 8 complexes have been performed for 30ns depicting RMSD plot with stabilized fluctuations, indicating the stability of the SOCS1-Peptide 8 complex.

### F. Systems driven synthetic biological circuit design

For synthetic circuits, one can emulate natural designs and/or use intuition and mathematical modeling to guide network choice. In both the cases, these approaches start from a single network – either based on some understanding of the mechanism, or on some intuition of the researcher. For addressing the present issue, we have designed mammalian tunable synthetic repressilator. The designed system contains *LacI, peptide 8, gfp* genes arranged in modular fashion under the influence of CMV promoter **(S4)**. IL6 synthetic gene circuit is constructed by assembling all necessary parts and pools. Parts are well defined DNA sequences having function in transcription or translation, whereas pools are the abstract places where free molecules of signal carriers are stored. Parts are composed into higher modules, the transcription units that interact by exchanging molecules of signal carriers such as transcription factor and small RNAs. Thus, in a circuit scheme, pools of signal carriers are the graphical interfaces among transcription units. It is worth noticing that the content of any pool plays a non-negligible role in determining the circuit dynamics. Parts and pools are modeled independently according to mass action kinetics by exchanging fluxes of signal carriers. These fluxes furthermore determine input/output in the genetic circuit and influence circuit performance. Parts that host interactions with signal carriers such as the promoter that binds RNA polymerase have access to the value of the concentration of free signal carrier molecules into their corresponding pools. The system is auto negative regulatory in nature due to the presence of Lac repressor gene (LacI) and its function is inhibited by IPTG (Fig6a). In absence of IPTG the system remains turn off (OFF STAGE) signifies no production of GFP (green fluorescent protein) and peptide 8, whereas in presence of IPTG the system is in ON STAGE representing the production of GFP and peptide 8 (Fig 6f). Both the stages have been confirmed through BoolNet package and transfected successfully in macrophage derived cell line (Fig 6h and 6i) **(S5)**. The simulation have been performed for 100 time units with graph representing oscillatory behavior confirming the auto negative regulatory nature of the designed system. The wiring graph obtained, shows that Lac repressor is the center/master for regulation of the whole system (Fig 6e). Using its time series data, convergence of statistical variables have been obtained, signifying modularity as well as orthogonality (Fig 6d). The null-cline point obtained through ODE solver states that the system has one stable state at a given time point, system will either be in an ON state or in OFF stage (Fig 6d). The results further imply that the system has tendency to follow same trajectory even in presence of external perturbations, known as canalization. *In silico* method to study circuit canalization is very similar to sensitivity analysis.

**Fig. 6.**
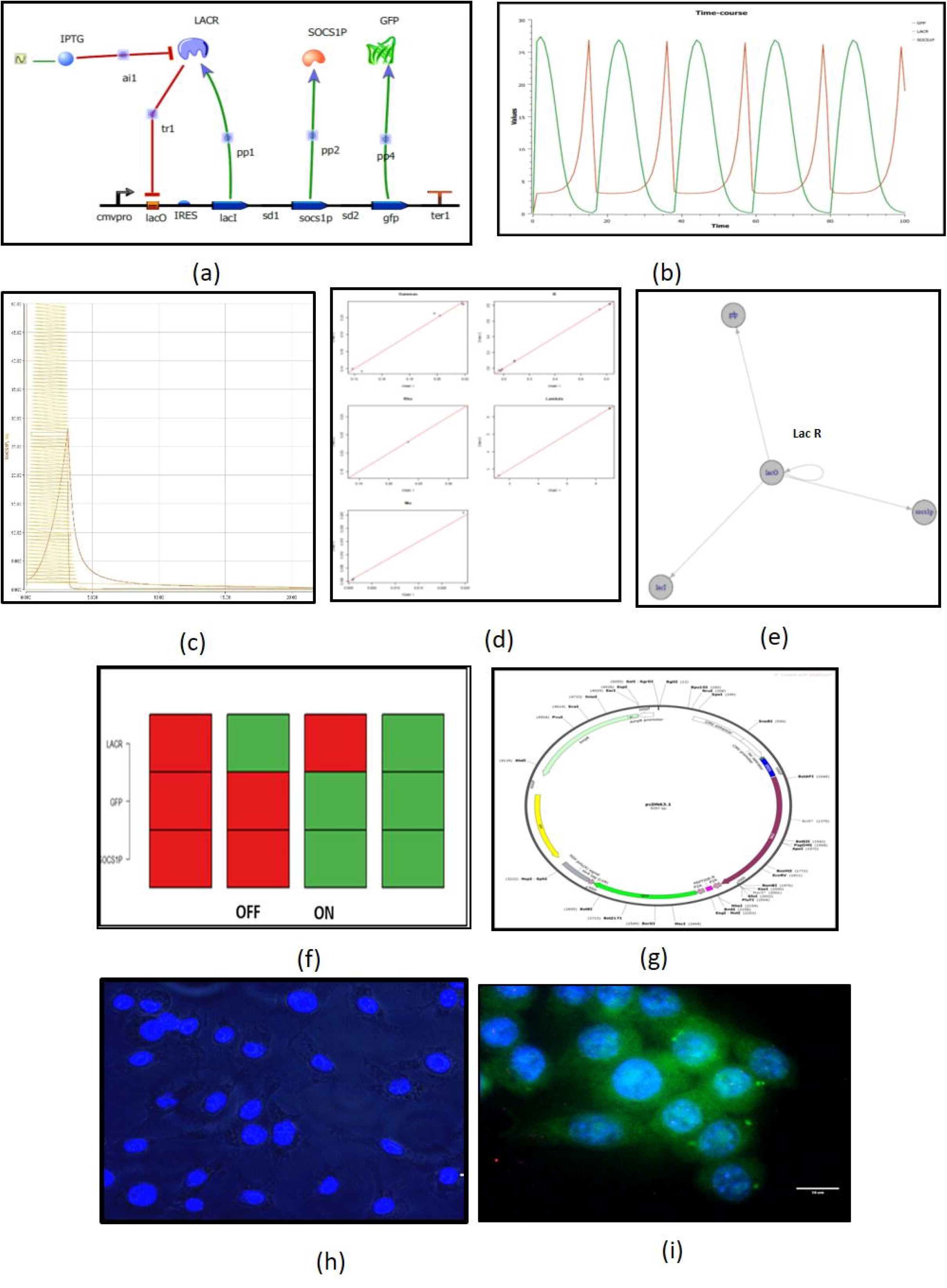
Synthetic circuit Design: (a) Modular arrangements of biological parts to receive an output in Tinker Cell (b) After simulating the system, the output is received in the form of oscillatory behavior of LACR, Peptide8 and GFP. (c) Nullcline form of synthetic circuit with states depicting synthetic circuit reaching its equilibrium state. (d) Convergence of statistical variables signifies the stability of the system. (e) Wiring of the circuit signifies the major regulatory axis as Lac Repressor gene. (f) Attractor states of synthetic circuits shows its ON and OFF stage (g) Plasmid map of designed synthetic circuit. Plasmid was transfected and expressed in RAW264.7 cell line, through IPTG induction, followed by DAPI staining showing (h) OFF stage of the system *in vitro* (no IPTG induction) (i) ON stage of the system *in vitro* (with IPTG induction).

#### *In vitro* validation

##### A. Cytokine profiling

To determine the efficacy of designed synthetic circuit, cytokine profiling for various groups were performed using Taqman chemistry. Miltefosine is taken as positive control and the study was divided into five groups namely Control (C), Infection(I), Empty Vector (EV), Transfection + Infection (CTI), Transfection + Infection + Miltefosine (CTIM).

During the initial interaction, cytokines having pro-inflammatory behavior are found to increase with the introduction and expression of synthetic circuit. The expression was further increased with the treatment of Miltefosine. If we quickly observe the cytokine profiling of the infectious state, TNF α shows a constant 2-3 fold change which symbolizes its role in parasite clearance during initial infection but the fold change of IL12 (2-5 fold change) is low as compared to fold change of IL10 (7-8 fold change). This predominantly shows the negative regulation of IL10 over IL12 which turns the polarization of macrophages towards M2 phenotype (Fig 6). Further adding to this, there was no expression of IFN γ and iNOS which is due to the constant increase in fold change of TGF beta (0-45 minute post infection), depicts that TGF β is strong anti-inflammatory cytokine which suppresses the expression of IFN γ and iNOS (Fig 7b & 7c). IL1β has both pro and anti-inflammatory action, the increase in fold change of IL1β in synchrony with IL10 and TGF β shows its predominant anti-inflammatory action in establishing infection during early state (Fig 7a & 7c). There was no expression of IL4 observed post one hour of infection.

**Fig.7.**
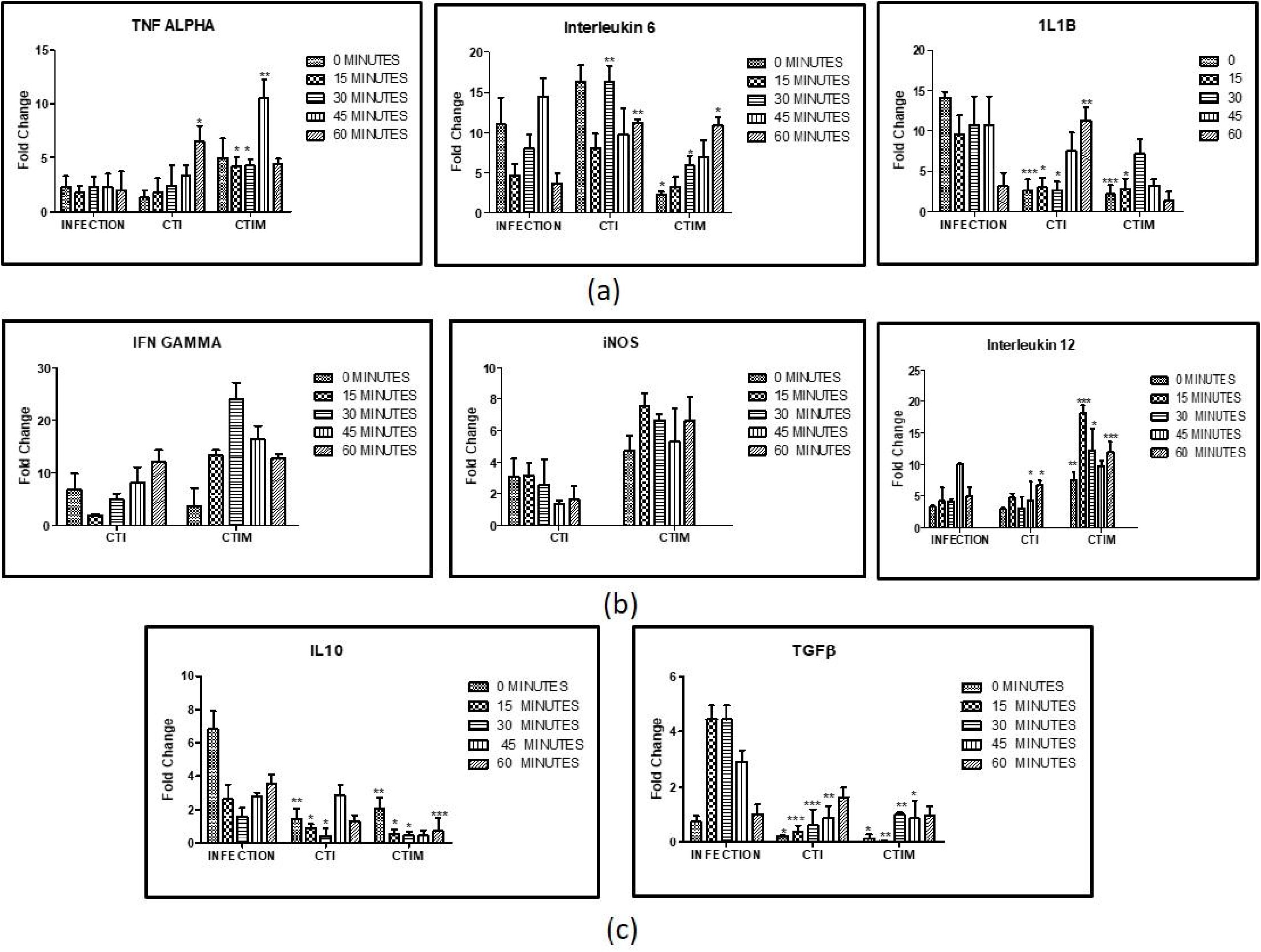
Cytokine profiling of *in vitro* validation of synthetic circuit: Macrophages were first transfected with synthetic circuit using, Polyethylenimine (3:1 ratio PEI: DNA) and then induced with 1mM IPTG for 48 hrs followed by infection with stationary phase promastigotes in 1:10 macrophage to parasite ratio for 24 hrs. Un-internalized parasites were washed off and samples were collected at different time points. (a) Cytokines released during initial interaction (b) Cytokines associated with M1 phenotype. The levels of IFN γ and iNOS has not been observed in Infection group therefore, bars are not shown. (c) Cytokines associated with M2 phenotype. The experiments were performed thrice. The error bars represents mean ± SD with p value *p < 0.05, **p < 0.01, ***p < 0.001

On introduction of designed synthetic circuit in infected cell, there is rapid increase of TNF α which has been observed with nearly 10 fold change (60 minutes post infection) and which further increase with Miltefosine treatment (Fig 7a). This proves that the designed circuit promotes the levels of TNF α in micro-environment establishing the anti-leishmanial response during early stage of infection. Furthermore, sharp fall in fold change of IL10 and TGF β have been observed. The fold change levels of IL10 have reached to 1-3 times from 7-8 times whereas the fold change of TGF β has been dropped from 4-5 times to 1-2 times (Fig 7c) representing the negative regulation of TNF alpha over IL10 and TGF beta and thus shifting the polarization towards M1. Although there is not much increase in fold change of IL12, but if we observe minutely, reciprocity in regulation have been observed between IL10 and IL12 at 45-60 min post infection (Fig 7a & 7c). The major increase in fold change of IFNγ and inducible nitric oxide synthase (iNOS) shows that introduction of designed synthetic circuit is tilting the macrophage phenotype towards classical activation, resulting in killing of parasite inside macrophages (Fig 7b).

##### B. Nitrite Estimation

The potential of the designed synthetic circuit was further validated by quantifying nitrite in the system, which is indicative of macrophage polarization towards M1 phenotype. The study is further divided into five groups as mentioned above and Lipophosphosaccharide (LPS) is taken as positive control. Production of nitrite among control, infection and empty vector are found to be similar but when designed synthetic circuit is induced in infected cells, nitrite production is increased which shows that the IL6 synthetic biological circuit is shifting macrophage polarization towards M1 phenotype. With the treatment of miltefosine, nitrate production is further increased (Fig 8c).

**Fig.8.**
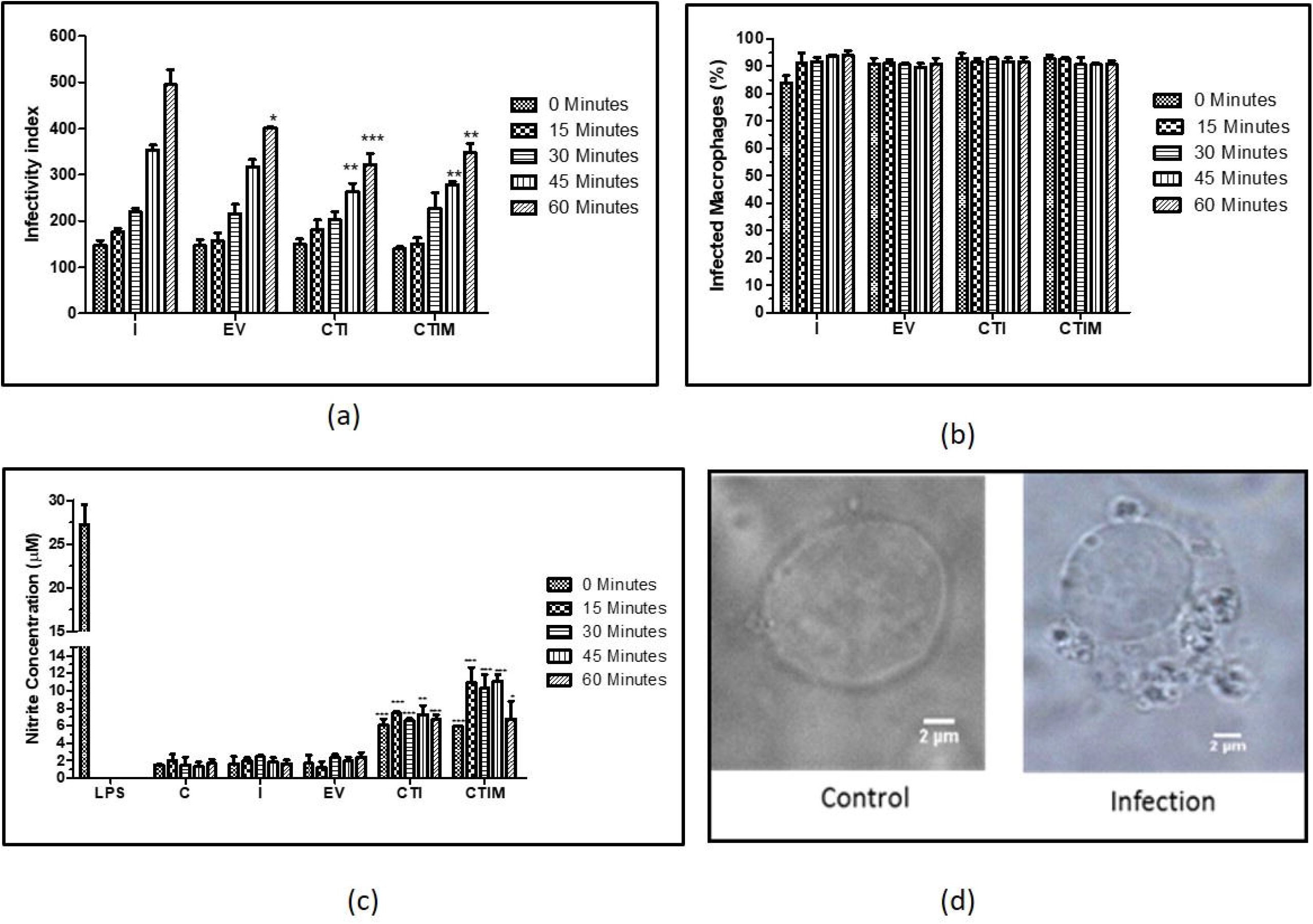
*In vitro* validation of designed synthetic circuit: (a) Estimation of parasite burden in terms of infectivity index in various infected and transfected groups. (b) The percentage of macrophage infected with *L. major* parasite (c) Nitrite estimation for various groups post one hour of infection in comparison with LPS treatment. (d) Transmitted microscopic image (100X) of control and infected RAW264.7 macrophage. The experiments were performed three times. One way ANOVA (with Tukey’s correction) have been used to perform statistical analysis of Infectivity index and student t test have been used to analyse Nitric oxide estimation The error bars represents mean ± SD with p value *p < 0.05, **p < 0.01, ***p < 0.001.

##### C. Parasite load assay

After nitrite estimation, parasite burden have been estimated in various groups (Fig.8a &8b). No change in parasite burden has been observed post 30 minutes of infection either in transfected or non-transfected system. During 45-60 minutes post infection, the fold change in iNOS and IFNγ (fig 7b) have been increased together with high nitrite production (Fig 8c) resulting in significant decrease in parasite load in transfected system (CTI and CTIM).

## Discussion

IL6 gene expression systems, designed in this study, deal with stochasticity due to the random nature of cellular dynamics associated. The effect of system non-linearity and stochasticity combined with global sensitivity analysis has given sufficient impetus for IL6 synthetic circuit modular analysis which in turn provides framework of retroactivity, all together on a system wide level. The model combined with experimental data captured the host-immune dynamics of parasitic infection and helped identify key components that is crucial for explaining individual variability of different cytokines for a dynamical cellular response. Identification of key components in these complex networks and linking these multivariate interactions to events at different physiological scales, for example, tissue level behavior that directly contributes to disease states, is the crux in systems immunology. During the process of model refinement in this paper, we have identified the ratio of SOCS1:SOCS3as 3:2 for establishment of infection which is further exploited as a target for designed therapeutics. The elevated levels of SOCS1 protein (60,000 molecules/ cell) have been mathematically quantified and found to inhibit the signaling of proinflammatory cytokine such as IL12, IFN γ, TNFα. Further this inhibition resulted in increased production of anti-inflammatory cytokines (around 2-7 fold change have been found as compared with control samples). Model analysis at various levels flux, sensitivity and principle component analysis represented the key reactions governing the dynamics of diseased state and SOCS1 playing a crucial role in the same. To add to this, structural analysis of these proteins helped identify specific regions responsible for its inhibitory action. The region is then targeted by designing set of peptides against it. The *in silico* design and analysis of the SOCS1-peptide complex ensures us to test the efficiency in *in vitro* condition. In order to make the delivery of the peptide more specific and less expensive, we opt for synthetic biology approach, wherein, an inducible gene regulatory circuit delivers the designed peptide at specific location (in present work it is in cytoplasm where there is production of SOCS1 protein). Here, the circuit design is precise as well as simple to avoid unnecessary complications during its transfection or else like a Rube Goldberg machine the designed circuit may look exciting *in silico* but would rarely yield informative results in wet lab conditions. The confirmatory analysis of the designed therapeutic shows a remarkable upregulation in pro-inflammatory cytokines such as TNF α, iNOS, IFN γ and IL12 and apparently down regulation of anti-inflammatory cytokines IL10 and TGF β. The present data is indicative of the effective functioning of the synthetic circuit. The results have further been confirmed through estimation of nitric oxide as well as parasite load in macrophages.

One of the major aspects of this current concept is by using system driven synthetic immunology approach, we have actually targeted the host system (which contribute to disease progression) rather than the parasite itself counteracting the issue of resistance development. The inducible nature of the synthetic circuit ensures the control over the designed product and its action over the host system at cellular level. Robustness/homeostasis of the synthetic circuit corresponds to the circuit capability to stabilize a quantity (e.g., the fluorescence level) against deviations from a given value (the one at steady state, for inference). Homeostasis attained via a designed structural motif/peptide inserted into the circuit helped to establish a temporal program of gene expression. The comparison of the response time at different steps quantified the delay in the output production due to the cascade length, made by a series/row of genes, the first regulating the second one, the second the third one and so on. Each regulation adding a step to the working gene cascade and is not easy to engineer *in vivo* because the noise in the output is amplified at intermediate fluorescence values for high cascade length. This might prevent synchronization of cell response over a population. Moreover, a careful fine-tuning of kinetics is necessary in order to assure proper signal propagation along the IL6 cascade. A negative feedback loop accelerates the response time of IL6 circuit and stabilizes protein concentration. As discussed, the major limitation of the systems comes when the therapy is taken at complex biological level such as tissue/organ or entire organism. At higher biological level, the system design is achieved with respect to the complexity of the biochemical network vis-a-vis combining the present design of the synthetic circuit with CRISPR-Cas9 system for its better performance in *in vivo* system (20). Nonetheless, it will make the system much more bulky which eventually affects its transfection efficiency. Fine tuning of the biological response may pose hindrance at higher level. Nevertheless, it is evident that synthetic biology approach is still among the prominent and most appropriate choice for designing new therapeutics regime because of its specificity, cost efficiency and less off target effects.

## MATERIALS AND METHODOLOGY

### In silico

#### A. Reconstruction and Analysis of IL6 Mathematical Model

Data fitting was performed with respect to the ratio of SOCS1/ SOCS3, obtained from wet lab experimentation (Western Blotting) followed by optimization wherein respective parametric changes have been performed to fine tune the models. The entire data sets were simulated with deterministic approach using 15s ODE solver followed by sensitivity analysis which quantifies the dependency of model trajectories upon variation in introduced parameters. Further, quantification of prediction uncertainties has been performed. Model construction, refinement and analysis have been performed in Simbiology Matlab Tool box (v7.11.1.866) and Copasi (v4.19).

#### B. Multi Objective Optimization and Evolvability

The macrophage phenotype network have been optimized by defining two objective function, f (1): SOCS1 associated with anti-inflammatory response, f(2): SOCS3 associated with pro-inflammatory response. We have used Multi-objective genetic algorithm (MOGA) to optimize the ratio of SOCS1:SOCS3(21). The optimization was performed in MATLABs’ Optimization toolbox (7.11.1.866) (MathWorks Inc.) using the function solver “gamultiobj”.

#### C. Target Identification and Protein-Protein Docking

The refined models have further been analyzed and reduced which crucified Suppressor of Cytokine Signaling 1 (SOCS1) as the target. 3D structure prediction have been done for various proteins of IL6 signaling complex (mSOCS1, mgp130, mIL6R and mJAK) using *ab initio* modeling techniques (Robetta) and homology modeling (Modeller 9.18) apart from mIL6 (PDB ID: 2L3Y). SOCS1 protein was docked with IL6 signaling complex to identify interfacial residues involved in interaction (Fig.5a). Most of the residues belong to SH2 domain of SOCS1 and henceforth non-conserved region around SH2 domain is targeted for peptide designing using Dead End Elimination algorithm. The non-conserved regions were identified through multiple sequence alignment of all the SOCS1 protein of mouse (MultAlign) (Fig.5c).

#### D. Peptide Design, Docking and MD Simulations

Peptide library was designed using deterministic search method (Dead End Elimination) and secondary structure was obtained through PEPstrMOD(22), followed by docking against SOCS1 through Autodock Vina (v1.5.6)(23) and interaction studies through Ligplot(24). Selected peptide-protein complex was further subjected to molecular dynamic simulations for 30ns to study its stability in physiological condition. MD simulation was performed using DESMOND 3.2 (D.E. Shaw Research) from Maestro 8.2 (25), in explicit TIP3P water model using orthorhombic box with a default 10nm cutoff PBC (periodic boundary condition) for a time period of 50ns with the time steps of 2 fs. The RMSD, RMSF and the trajectories were analyzed using simulation event analysis in Desmond 3.2.

#### E. Synthetic Circuit Design and Quasipotential Landscape

Designing and simulation of synthetic circuit was performed in Tinker cell (v1.2.693) (26). Modularity and orthogonality of the circuit was confirmed through BoolNet (27). Time series data for 100 time units were generated through Gene Regulatory Network Inference using Time Series (GRENITS) (v 1.24.0)(28, 29). Various parts of synthetic circuit were obtained from Registry of Standard Biological Parts (iGEM) (Table 2) and assembled in Snapgene (v3.2.1), followed by its procurement in the form of plasmid from Gene art Thermofisher Scientific. Stability of the synthetic circuit was achieved by obtaining its nullcline state through Berkeley Madonna (Version 9.1.3), followed by obtaining quasipotential landscape through equation Vq=−((LacR)+(peptide:8))*DT derived from the Waddington’s epigenetic landscape (30).

**Table 2.** Registry of standard biological parts with parts registry number **(Sequences of the same enlisted in S7)**

### In vitro

#### A. Cell culture and parasites

The pathogenic promastigote form of *Leishmania major* strain (MHOM/Su73/5ASKH) were maintained in Roswell Park Memorial Institute (RPMI) 1640 with 20 % fetal bovine serum (Sigma) and 50 U/ml penicillin. The parasite was passaged regularly through BALB/c by injecting stationary phase promastigotes in subcutaneous region in order to maintain its virulence (31). The murine macrophage cell line RAW264.7 was maintained at 37 °C with 5% CO_2_ in Dulbecco’s Modified Eagle Medium (DMEM) with 10% fetal bovine serum and penicillin (100ug/ml).

#### B. Reagents, Antibodies, Probes and Constructs

All other chemicals were from Sigma-Aldrich, unless indicated otherwise. Antibodies for Western blotting such as anti-IL6 (be006) from Biocell, anti-phospho STAT3 (S2690), from Sigma and mouse IL6 (#5210), anti-phospho STAT1 (#9177), anti-SOCS1 (#2923) and anti-SOCS3 (#3950) were obtained from Cell Signaling Technology (CST). Taqman Chemistry was used to perform and quantify cytokine expression levels in samples. Mouse specific taqman probes (4331182) and Mastermix (4304437) were obtained from Thermofisher Scientific. Peptidoglycan from *Staphylococcus aureus* (PGN-SA) (#tlrl-pgns2) and Mab-mTLR2 (#mab-mtlr2) from InvivoGEN were procured. Designed synthetic circuit was procured in the form of plasmid from GeneArt, Thermofisher Scientific.

#### C. Macrophage and Parasite infection

For *in vitro* experimentation, RAW 264.7 cell line was infected with stationary phase promastigotes in 1:10 macrophage/ parasite ratio for 24 hrs, followed by washing of un-internalized parasite and incubating the infected cells in DMEM with 10% FBS.

#### D. Transfection of macrophages

Macrophages were transfected with designed synthetic construct (plasmid form) using Polyethylenimine transfection reagent in a 3:1 ratio of PEI to DNA (w/w). The transfected cells were induced by 1mM IPTG (Isopropyl β-d-1-thiogalactopyranoside) for 48 hrs followed by infection. Transfected cells were visualized for GFP expression on EVOS FL fluorescence microscope.

#### E. mRNA isolation, RT PCR and Real time PCR

For cytokine profiling, after washing un-internalized parasite, cells were lysed and total RNA was isolated at 0 min, 15min, 30 min, 45 min and 60 min post-infection. The total RNA was isolated using TRI Reagent as per the manufacturer’s instructions. The cDNA synthesis was done using 2ug of total RNA through high Capacity cDNA kit (Invitrogen) as per the manufacturer’s instructions.

Q-PCR was performed on StepOnePlus Real-Time PCR System (Thermo Scientific). For each reaction, 5ul Taqman Master mix (Invitrogen), 1ug cDNA as Template, 0.5 ug Taqman probes **(S6)** was taken, and the reactions were performed on thin-wall 0.1 ml fast 96 well plate (Applied Biosystems) for a total of 10 ul reaction mix. Relative quantitation was done using the comparative threshold (ΔΔCT) method. The mRNA expression levels of the target genes were normalized against those of β actin levels and expressed as relative fold change compared with untreated controls.

#### F. Western Blotting

##### Cross talk validation

For activation, RAW 264.7 cell line was treated with TLR2 activator PGN-SA (Peptidoglycan of *Saccharomyces aureus*) for 24 hours and for inhibition, culture was treated with 2ug/ml of mTLR2 before activation. The activation of IL6 pathway has been performed by treating cell with mIL6 (50ng/ml) for 24hrs and for inhibition, anti-IL6 antibody (1ug/ml) treatment was given for 1 hour prior IL6 treatment.

##### SOCS1/SOCS3 validation

The macrophage derived cell line RAW 264.7 cell line were infected with promastigote form of parasite in 1: 10 ratio, followed by 24hrs incubation and removal of undigested parasite. The culture was then treated with 50ng/ml of mouse IL6 protein for another 1 hour followed by sample collection.

For Western blotting, cells were treated with indicated reagents and lysed with lysis buffer (50 mM Tris [pH 7.5], 250 mM NaCl, 50mM NaF, 10% glycerol, 5 mM EDTA, 0.5 mM Sodium orthovanadate, and 0.5% TritonX), and a protease inhibitor mixture, by incubation on ice for 20 min followed by centrifugation of lysates (15,000 rpm, at 4°C for 20 min), and supernatants were quantified by Bicinchoninic acid kit (Thermofisher scientific). Equal amount of protein was loaded on SDS-PAGE, and resolved proteins were transferred to nitrocellulose membrane (Millipore, Billerica, MA) and blocked with 3% Bovine serum albumin in TBST (20 mM Tris [pH 7.5], 150 mM NaCl, and 0.1% Tween 20). Membranes were incubated with primary Antibody (1:1000 dilution) at 4°C overnight, followed by washing with TBST, and incubated with HRP-conjugated secondary Ab. Immuno-reactive bands were visualized with the Luminol reagent (Santa Cruz Biotechnology, Santa Cruz, CA). Densitometric analysis of bands was performed using Image J software.

#### G. Parasite load assay

After specific treatment, macrophages were washed with 1X cold PBS followed by fixation in 4% paraformaldehyde (PFA), and permeabilization in 0.1% Triton X. Then the cells were stained with DAPI (1μg/ml). Parasites per macrophage were calculated using EVOS FL fluorescence microscope and they were presented in terms of infectivity index (percentage of infected cells x number of parasites per infected cells).

#### H. Estimation of ROS production

Presence of Nitrite in culture media is an indicator of Nitric oxide production by cells (precisely macrophage polarization towards M1 phenotype). The Cell culture supernatant in 150 μl volume is treated with 20 μl of Griess reagent (0.1% N-(1-naphthyl)ethylenediamine and 1% sulfanilic acid in equal volume; Thermofisher Scientific) to set a total volume of 300 μl per reaction and incubated for 10 minutes at room temperature followed by colorimetric estimation at 540nm on.

#### I. Animal Maintenance

Female BALB/c mice, 6–8 weeks old with 18–20 g weight, originally procured from The Jackson Laboratory (Bar Harbor, ME) and maintained in the Experimental Animal Facility of National Centre for Cell Science (NCCS), Pune. Animals were used according to the Institutional Animal Ethical Committee–approved animal use protocol (IAEC Project Number-IAEC/2016/B-269).

#### J. Statistical analysis

The *in*-*vitro* experiments were performed in triplicates. The error bars are represented as mean□±□s.d. The statistical significance between the indicated experimental and control groups was deduced by using Student’s *t*-test and One way ANOVA (with Tukey’s correction).

## Supporting information

Concentration of components of mathematical model

S2A-S2B: S2A: Peptide sequences and Docking Score S2B: Characteristic of Peptide8 obtained from ExPASY ProtParam tool

S3A-S3B: S3A: RMSF Plot of SOCS1-P8 Complex S3B: Physical parameters during 30ns MD simulation

DNA sequence of parts used in the synthetic circuit

Insert Verification

Raw Images of Western Blots

## Abbreviations

AIF: Anti-inflammatory factor
CMV: Cytomegalovirus
CT: Comparative threshold
CTI: Control + Transfected + Infected
CTIM: Control + Transfected + Infected + Miltefosine
DAPI: 4′,6-diamidino-2-phenylindole
DMEM: Dulbecco’s Modified Eagle Medium
DSM: Diseased State Model
EDTA: Ethylenediaminetetraacetic acid
EV: Empty Vector
GFP: Green Fluorescent Protein
HSM: Healthy State Model
IFNγ: Interferon gamma
IPTG: Isopropyl β- d-1-thiogalactopyranoside
iGEM: International Genetically Engineered Machine
iNOS: Inducible nitric oxide synthase
IL6: Interleukin 6
IL1β: Interleukin 1 beta
IL10: Interleukin 10
JAK/STAT: Janus kinase/signal transducer and activator of transcription
LACR: Lactose Repressor
LPS: Lipopolysaccharide
MOGA: Multi-objective genetic algorithm
ODE: Ordinary differential equation
PBC: Periodic boundary condition
PCA: Principal component analysis
PFA: Paraformaldehyde
PGN-SA: Peptidoglycan from *Staphylococcus aureus*
PEI: Polyethylenimine
RPMI: Roswell Park Memorial Institute
RMSD: Root-mean-square deviation
RMSF: Root mean square fluctuation
SH2: Src Homology 2
SOCS: Suppressor Of Cytokine Signaling
TIP3P: Transferable intermolecular potential with 3-point
TLR: Toll like Receptors

## Supplemental Material

**S1:** Concentration of components of mathematical model

**S2A-S2B:**

S2A: Peptide sequences and Docking Score

S2B: Characteristic of Peptide8 obtained from ExPASY ProtParam tool

**S3A-S3B:**

S3A: RMSF Plot of SOCS1-P8 Complex

S3B: Physical parameters during 30ns MD simulation

**S4:** DNA sequence of parts used in the synthetic circuit

**S5**: Insert Verification details

**S6:** Invitrogen® Assay ID for RT PCR probes.

**S7:** Raw Images of Western Blots

## Acknowledgements

Bhavnita Soni acknowledges her Senior Research fellowship from INSPIRE, Department of Science and Technology (DST), Ministry of Science and Technology, Government of India. Authors would like to thank Cell Repository and Media Section of National Centre for Cell Science (NCCS). We also extend our thanks to the Director, NCCS for supporting Bioinformatics and High Performance Computing Facility (BHPCF) at NCCS, Pune, India.

## REFERENCES

1. Croker BA, Kiu H, Nicholson SE. 2008. SOCS regulation of the JAK/STAT signalling pathway. Semin Cell Dev Biol.

2. Duncan SA, Baganizi DR, Sahu R, Singh SR, Dennis VA. 2017. SOCS proteins as regulators of inflammatory responses induced by bacterial infections: A review. Front Microbiol 8:1–15.

3. Sun K, Salmon S, Yajjala VK, Bauer C, Metzger DW. 2014. Expression of Suppressor of Cytokine Signaling 1 (SOCS1) Impairs Viral Clearance and Exacerbates Lung Injury during Influenza Infection. PLoS Pathog.

4. Chien H, Alston CI, Dix RD. 2018. Suppressor of Cytokine Signaling 1 (SOCS1) and SOCS3 Are Stimulated within the Eye during Experimental Murine Cytomegalovirus Retinitis in Mice with Retrovirus-Induced Immunosuppression. J Virol 92.

5. Chandrakar P, Parmar N, Descoteaux A, Kar S. 2020. Differential Induction of SOCS Isoforms by Leishmania donovani Impairs Macrophage–T Cell Cross-Talk and Host Defense. J Immunol.

6. Soni B, Saha B, Singh S. 2018. Systems cues governing IL6 signaling in leishmaniasis. Cytokine.

7. Yang L, Ebrahim A, Lloyd CJ, Saunders MA, Palsson BO. 2019. DynamicME: Dynamic simulation and refinement of integrated models of metabolism and protein expression. BMC Syst Biol.

8. Hahl SK, Kremling A. 2016. A comparison of deterministic and stochastic modeling approaches for biochemical reaction systems: On fixed points, means, and modes. Front Genet.

9. Shain KH, Yarde DN, Meads MB, Huang M, Jove R, Hazlehurst LA, Dalton WS. 2009. β1 integrin adhesion enhances IL-6-mediated STAT3 signaling in myeloma cells: Implications for microenvironment influence on tumor survival and proliferation. Cancer Res.

10. Fisher DT, Appenheimer MM, Evans SS. 2014. The two faces of IL-6 in the tumor microenvironment. Semin Immunol.

11. Pradhan AD, Manson JE, Rifai N, Buring JE, Ridker PM. 2001. C-reactive protein, interleukin 6, and risk of developing type 2 diabetes mellitus. J Am Med Assoc.

12. Smole A, Lainšček D, Bezeljak U, Horvat S, Jerala R. 2017. A Synthetic Mammalian Therapeutic Gene Circuit for Sensing and Suppressing Inflammation. Mol Ther.

13. Ye H, Xie M, Xue S, Hamri GC El, Yin J, Zulewski H, Fussenegger M. 2017. Self-adjusting synthetic gene circuit for correcting insulin resistance. Nat Biomed Eng.

14. Kis Z, Pereira HSA, Homma T, Pedrigi RM, Krams R. 2015. Mammalian synthetic biology: Emerging medical applications. J R Soc Interface.

15. Ye H, Fussenegger M. 2014. Synthetic therapeutic gene circuits in mammalian cells. FEBS Lett.

16. Wroblewska L, Kitada T, Endo K, Siciliano V, Stillo B, Saito H, Weiss R. 2015. Mammalian synthetic circuits with RNA binding proteins for RNA-only delivery. Nat Biotechnol.

17. Perry N, Ninfa AJ. 2012. Synthetic networks: Oscillators and toggle switches for escherichia coli. Methods Mol Biol.

18. Elowitz MB, Leibier S. 2000. A synthetic oscillatory network of transcriptional regulators. Nature.

19. de Veer MJ, Curtis JM, Baldwin TM, DiDonato JA, Sexton A, McConville MJ, Handman E, Schofield L. 2003. MyD88 is essential for clearance of Leishmania major: Possible role for lipophosphoglycan and Toll-like receptor 2 signaling. Eur J Immunol.

20. Xu X, Qi LS. 2019. A CRISPR–dCas Toolbox for Genetic Engineering and Synthetic Biology. J Mol Biol.

21. Deb K, Pratap A, Agarwal S, Meyarivan T. 2002. A fast and elitist multiobjective genetic algorithm: NSGA-II. IEEE Trans Evol Comput.

22. Singh S, Singh H, Tuknait A, Chaudhary K, Singh B, Kumaran S, Raghava GPS. 2015. PEPstrMOD: Structure prediction of peptides containing natural, non-natural and modified residues. Biol Direct.

23. Trott oleg, Arthur J. Olson. 2010. AutoDock Vina: Improving the Speed and Accuracy of Docking with a New Scoring Function, Efficient Optimization, and Multithreading. J Comput Chem.

24. Wallace AC, Laskowski RA, Thornton JM. 1995. Ligplot: A program to generate schematic diagrams of protein-ligand interactions. Protein Eng Des Sel.

25. Bowers KJ, Chow E, Xu H, Dror RO, Eastwood MP, Gregersen BA, Klepeis JL, Kolossvary I, Moraes MA, Sacerdoti FD, Salmon JK, Shan Y, Shaw DE. 2006. Scalable algorithms for molecular dynamics simulations on commodity clustersProceedings of the 2006 ACM/IEEE Conference on Supercomputing, SC’06.

26. Chandran D, Bergmann FT, Sauro HM. 2009. TinkerCell: Modular CAD tool for synthetic biology. J Biol Eng.

27. Müssel C, Hopfensitz M, Kestler HA. 2010. BoolNet-an R package for generation, reconstruction and analysis of Boolean networks. Bioinformatics.

28. Morrissey ER, Juárez MA, Denby KJ, Burroughs NJ, Ideker T. 2011. On reverse engineering of gene interaction networks using time course data with repeated measurementsBioinformatics.

29. Morrissey E. 2012. GRENITS: Gene Regulatory Network Inference Using Time Series. R package version 1.24.0. 1–5.

30. Bhattacharya S, Zhang Q, Andersen ME. 2011. A deterministic map of Waddington’s epigenetic landscape for cell fate specification. BMC Syst Biol.

31. Kébaïer C, Louzir H, Chenik M, Ben Salah A, Dellagi K. 2001. Heterogeneity of wild *Leishmania major* isolates in experimental murine pathogenicity and specific immune response. Infect Immun.

