## Supplementary material for "SOCS1/SOCS3 Immune Axis Modulates Synthetic Perturbations in IL6 Biological Circuit for Dynamical Cellular Response": Concentration of components of mathematical model

**S1**

| **COMPONENTS** | **HSM**  **(molecules)** | **DSM**  **(molecules)** |
| --- | --- | --- |
| TLR2 | 1000 | 1000 |
| TLR6/1 | 1000 | 1000 |
| IL6-IL6R | 1000 | 1000 |
| IL6R | 1000 | 1000 |
| Membrane IL6 | 1000 | 1000 |
| LPG | 100 | 1000 |
| IL6-IL6R-GP130 | 2000 | 2000 |
| IFN-g | 1000 | 200 |
| IFN-gR | 1000 | 150 |
| IFNg-IFNgR | 1000 | 150 |
| gp130 | 1000 | 1000 |
| TLR2/6-LPG | 900 | 900 |
| IL10 | 100 | 1000 |
| IL10R | 100 | 1000 |
| IL10-IL10R Complex | 10000 | 10000 |
| MyD88 | 800 | 800 |
| IRAK1-IRAK4 | 700 | 700 |
| TRAF6 | 600 | 600 |
| TAK1-TAB1/2 | 500 | 500 |
| IKKbeta | 400 | 400 |
| NFkB | 300 | 300 |
| Cytoplasm IL6 | 100 | 100 |
| STAT3.P | 0 | 10000 |
| STAT1.P | 80000 | 0 |
| JAK1 | 20000 | 20000 |
| 2.STAT1 | 40000 | 0 |
| 2.STAT3 | 0 | 9000 |
| JAK2/1 | 2000 | 2000 |
| SOCS1 | 10000 | 10000 |
| TLP2 | 200 | 200 |
| NFkB | 200 | 200 |
| Nucleus IL6 | 250 | 250 |
| Nucleus 2.STAT1 | 6000 | 0 |
| Nucleus 2.STAT3 | 0 | 30000 |
| Nucleus SOCS3 | 10000 | 10000 |
| Nucleus STAT1 | 100 | 100 |
| Nucleus STAT3 | 100 | 100 |
| Nucleus SOCS1 | 4000 | 50000 |
| iNOS | 0 | 0 |
| AIF | 0 | 0 |
