## Supplementary material for "SOCS1/SOCS3 Immune Axis Modulates Synthetic Perturbations in IL6 Biological Circuit for Dynamical Cellular Response": S2A-S2B: S2A: Peptide sequences and Docking Score S2B: Characteristic of Peptide8 obtained from ExPASY ProtParam tool

**S2A. Table of peptide sequences and Docking score**

| **S no.** | **Motif** | **Peptide**  **Nomenclature** | **Peptide** | **Docking score** |
| --- | --- | --- | --- | --- |
| 1. | APGDTHFRTFRSHS | P1 | VHAKNSVDNADNTN | -6.2 |
|  |  | P2 | AHVKNTAENVEQSQ | -6.0 |
|  |  | P3 | VHAKNSVDNADN | -5.1 |
|  |  | P4 | AHVKNTAENVEQ | -6.4 |
|  |  | P5 | ADNTNCDCADDL | -6.4 |
| 2 | GDTHFRTFRSHS | P6 | AKNSADNVDNSQ | -5.5 |
|  |  | P7 | VRQTVEQADQSN | -5.9 |
| 3 | **SHSDYRRTTRTSAL** | **P8** | **NSQKADDLVDNNVI** | **-5.9** |
|  |  | P9 | QTNKVDELAQNNAI | -6.5 |
|  |  | P10 | NSQKADDLVDNN | -6.0 |
|  |  | P11 | QTNKVDELAQNN | -5.2 |
| 4. | RIVAAVGRENLA | P12 | DLAVVAADKSIV | -5.7 |
|  |  | P13 | ELVAAAVEKTLG | -5.9 |
| 5. | VAAVGRENLARI | P14 | AVVAVDNSIVDL | -4.7 |
|  |  | P15 | AVVAAENTIVEL | -6.4 |

**S2B: Characteristic of Peptide8 obtained from ExPASY ProtParam tool**

| Number of amino acids: | 14 |
| --- | --- |
| Molecular weight: | 1544.64 Da |
| Theoretical pI: | 3.93 |

i. **Amino acid composition:**

| Ala (A) 1 7.1% | Arg (R) 0 0.0% |
| --- | --- |
| Asn (N) 3 21.4% | Asp (D) 3 21.4% |
| Cys (C) 0 0.0% | Gln (Q) 1 7.1% |
| Glu (E) 0 0.0% | Gly (G) 0 0.0% |
| His (H) 0 0.0% | Ile (I) 1 7.1% |
| Leu (L) 1 7.1% | Lys (K) 1 7.1% |
| Met (M) 0 0.0% | Phe (F) 0 0.0% |
| Pro (P) 0 0.0% | Ser (S) 1 7.1% |
| Thr (T) 0 0.0% | Trp (W) 0 0.0% |
| Tyr (Y) 0 0.0% | Val (V) 2 14.3% |
| Pyl (O) 0 0.0% | Sec (U) 0 0.0% |

ii. **Atomic composition:**

| **ATOM** | **ATOMIC WEIGHT** |
| --- | --- |
| Carbon C | 63 |
| Hydrogen H | 105 |
| Nitrogen N | 19 |
| Oxygen O | 26 |
| Sulfur S | 0 |

Formula: C_63_H_105_N_19_O_26_	Total number of atoms: 213

iii. **Instability index:**

The instability index (II) is computed to be 6.24. This classifies the protein as stable.

iv. **Aliphatic index:** 104.29
v. **Grand average of hydropathicity (GRAVY)**: -0.764
