## Supplementary material for "SOCS1/SOCS3 Immune Axis Modulates Synthetic Perturbations in IL6 Biological Circuit for Dynamical Cellular Response": S3A-S3B: S3A: RMSF Plot of SOCS1-P8 Complex S3B: Physical parameters during 30ns MD simulation

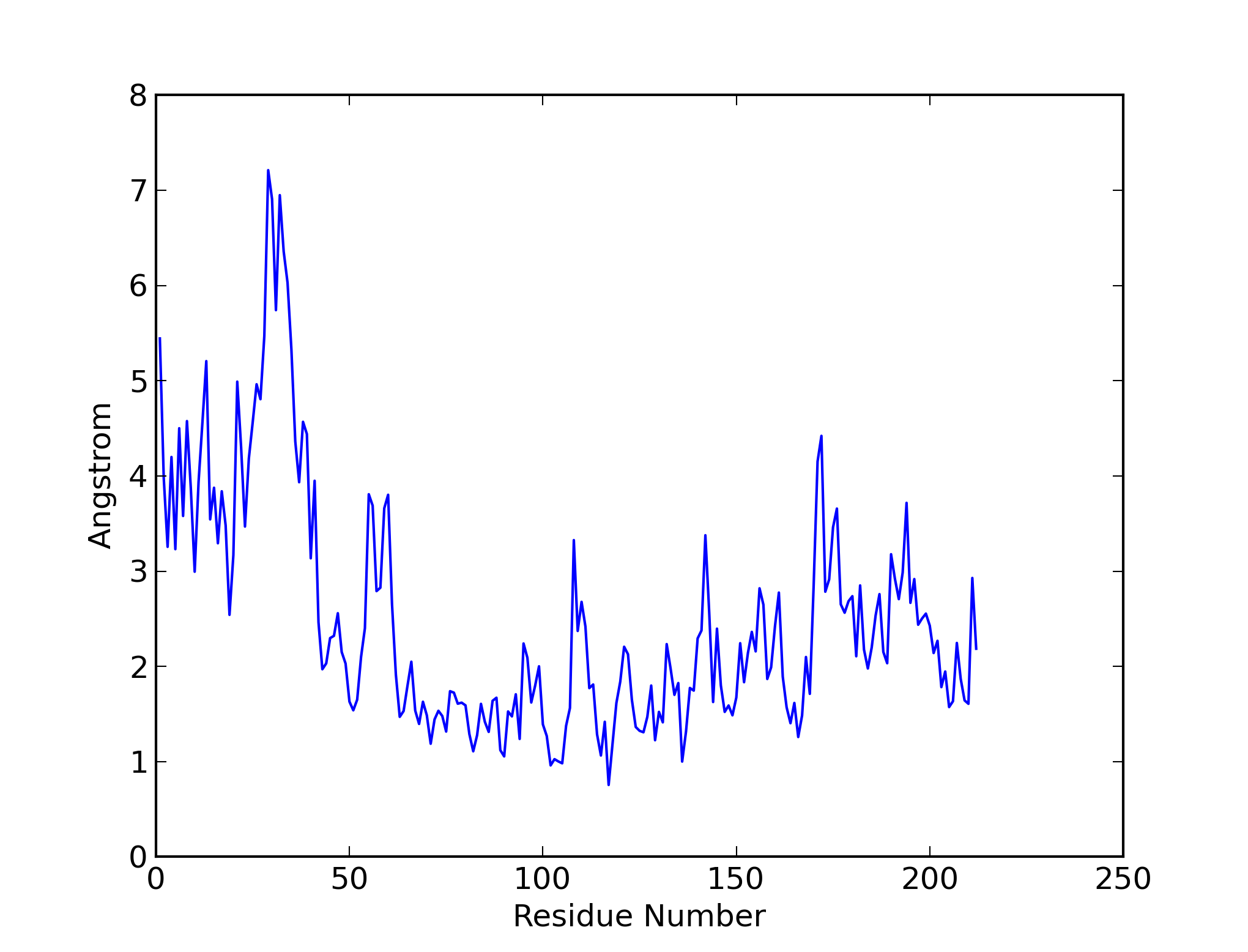

**RMSF plot for SOCS1-P8 complex**

**S3B : Physical parameters during 30ns MD simulation**
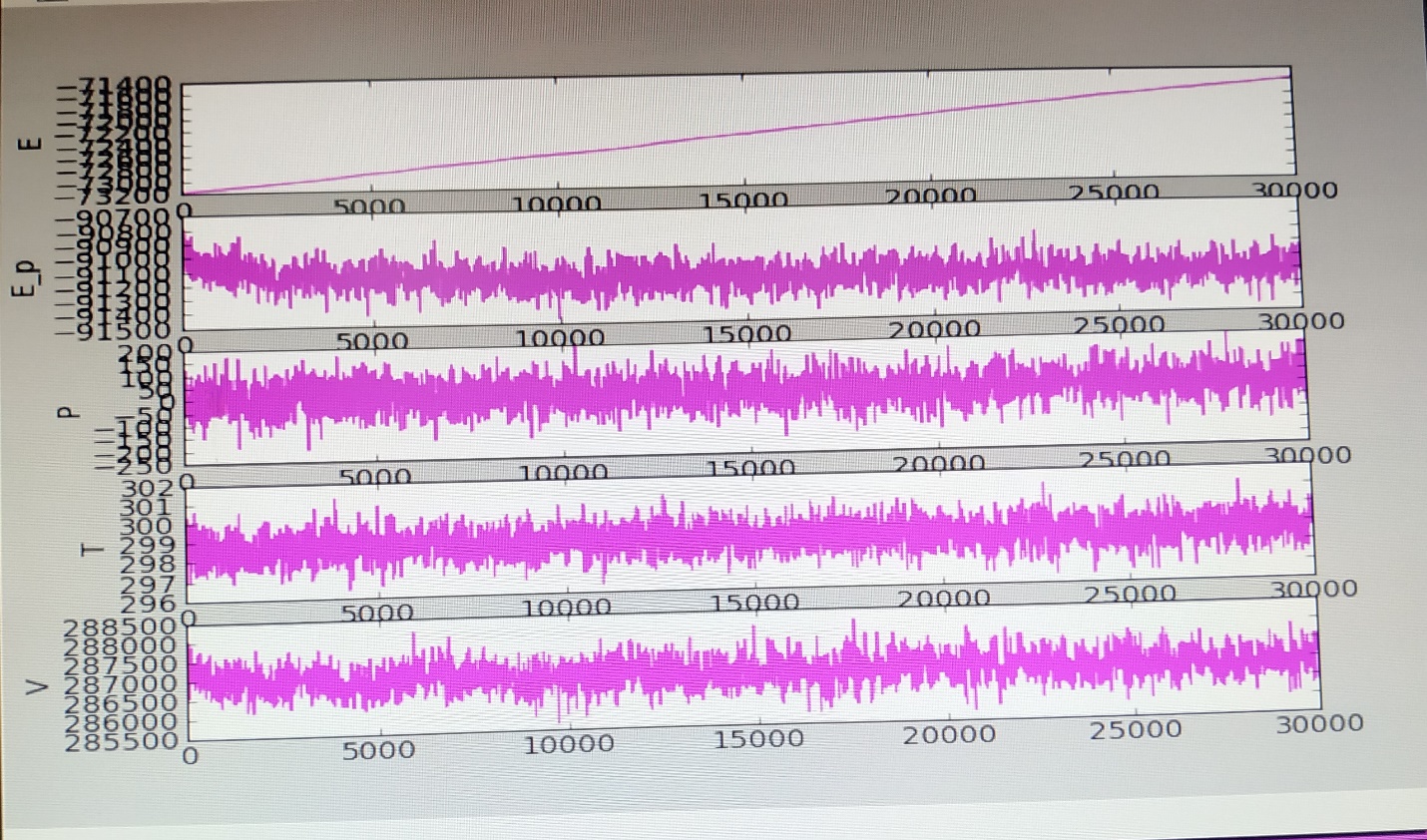
