## Supplementary material for "SOCS1/SOCS3 Immune Axis Modulates Synthetic Perturbations in IL6 Biological Circuit for Dynamical Cellular Response": Insert Verification

Printed on Aug 10, 2019
Sequence Scanner 1.0

### Quality Control Report

#### Contiguous Read Length

Number of traces (N) = 3
0 traces have short CRL.
0 trace have medium CRL.
2 traces have long CRL.
Mean = 797
Median = 796
Range = 733 - 860
Standard Deviation = 90

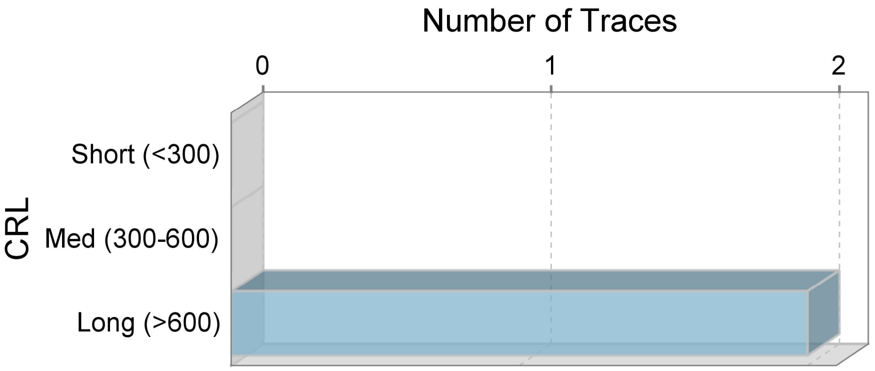


#### Legend

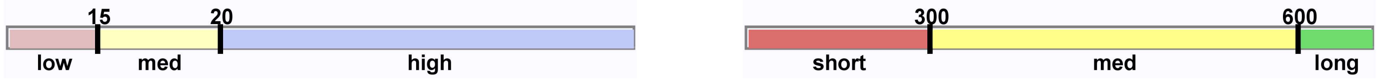


| Trace File Name | Well# | Cap# | Trace Score | CRL | QV20+ | Signal Intensity A | C | G | T | Comments |
|---|---|---|---|---|---|---|---|---|---|---|
| SAHAB_T7_F_A_H08.ab1 | H8 | 25 | 44 | 733 | 748 | 51 | 66 | 83 | 75 | |
| SAHAB_T7_F_F08.ab1 | F8 | 27 | 46 | 860 | 863 | 86 | 118 | 135 | 128 | |
| SAHAB_T7_R_G08.ab1 | G8 | 26 | N/A | N/A | N/A | 22 | 20 | 20 | 24 | |
