## Supplemental Data 1 for "SOCS1/SOCS3 Immune Axis Modulates Synthetic Perturbations in IL6 Biological Circuit for Dynamical Cellular Response"

**S6:**

|  | **Invitrogen® Assay ID** | **Gene** |
| --- | --- | --- |
| 1. | Mm00445259_m1 | IL4 |
| 2. | Mm00446190_m1 | IL6 |
| 3. | Mm01288386_m1 | IL10 |
| 4. | Mm00434165_m1 | IL12α |
| 5. | Mm00434228_m1 | IFN-γ |
| 6. | Mm00443258_m1 | TNFα |
| 7. | Mm02619580_g1 | ACT b |
| 8. | Mm01321739_m1 | TGFB |
| 9. | Mm00434228_m1 | IL1B |
