## Supplementary material for "SOCS1/SOCS3 Immune Axis Modulates Synthetic Perturbations in IL6 Biological Circuit for Dynamical Cellular Response": Raw Images of Western Blots

**S7**

**CROSS TALK POINTS:**

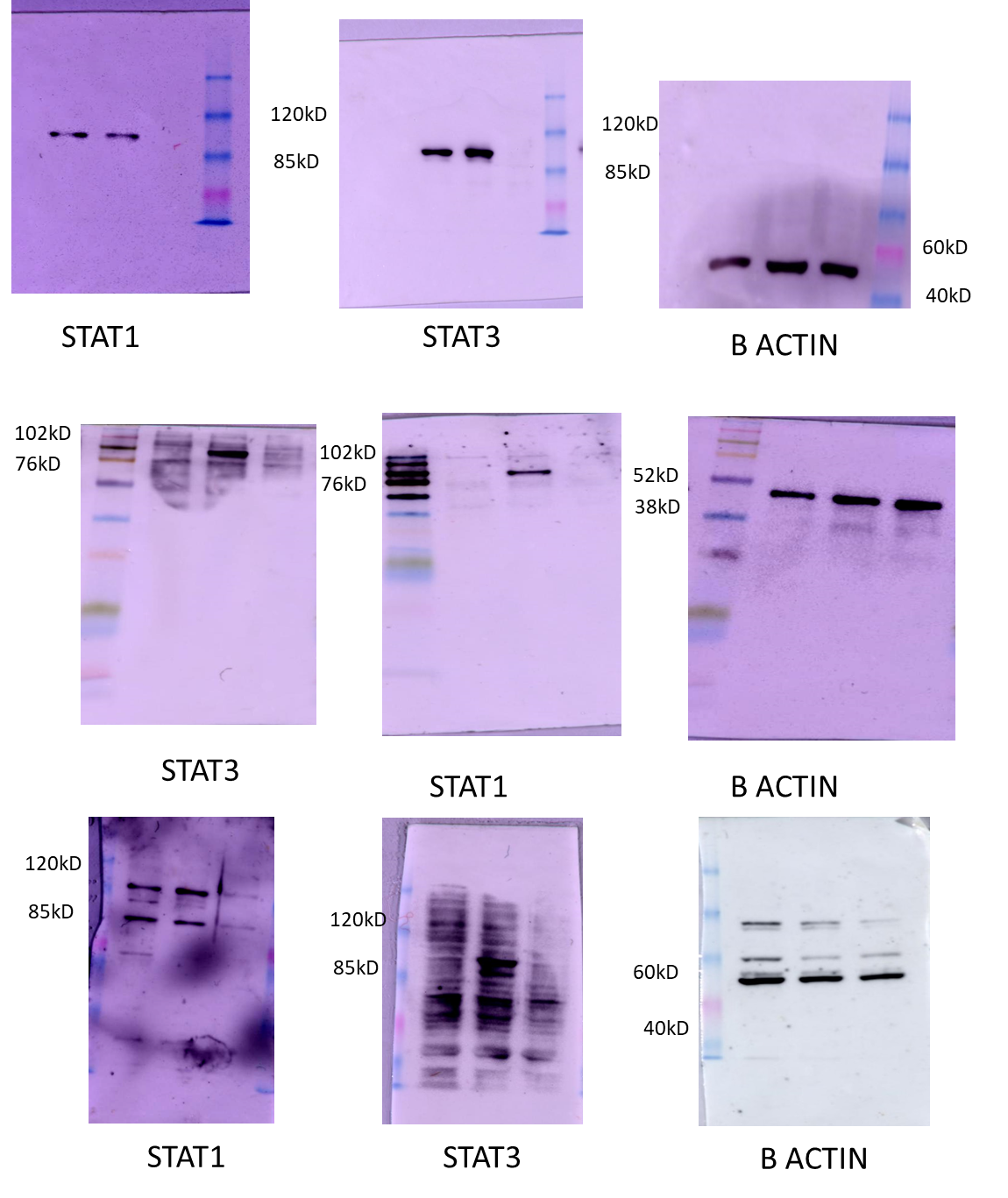

A. **Experimental validation of SOCS1:SOCS3 ratio:**

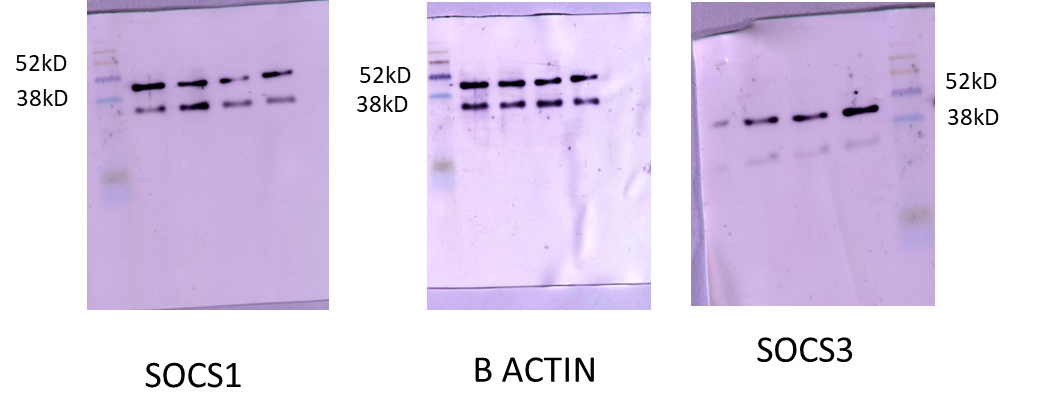

B. **Estimation of IL6 levels in control and infection:**

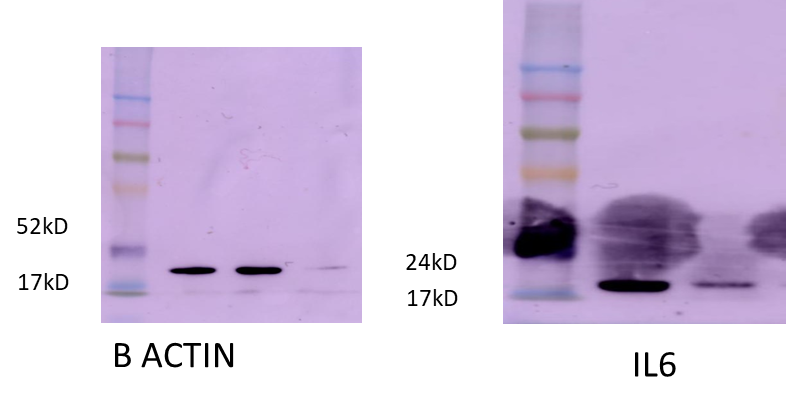
