## Supplementary material for "SOCS1/SOCS3 Immune Axis Modulates Synthetic Perturbations in IL6 Biological Circuit for Dynamical Cellular Response": Table1

**Table 1: Reaction that governs the disease progression at cellular level**

| **S.No.** | **Reactions** | **Flux**  **(molecules/sec)** | **PCA**  **Score** |
| --- | --- | --- | --- |
| 1. | TLR2/6-LPG -> MyD88 | 207.4748058 | 0.66 |
| 2. | JAK1 + STAT3{CYTOSOL} -> STAT3.P | 666.8136585 | 0.73 |
| 3. | SOCS1{NUCLEUS} -> SOCS1{CYTOSOL} | 15045.06 | 0.73 |
| 4. | SOCS3{NUCLEUS} -> SOCS3{CYTOSOL} | 4999.92 | 0.78 |
