## Supplementary material for "SOCS1/SOCS3 Immune Axis Modulates Synthetic Perturbations in IL6 Biological Circuit for Dynamical Cellular Response": Table 2

**Table 2.** Registry of standard biological parts with parts registry number **(Sequences of the same enlisted in S7)**

| **S.No.** | **Biological Parts** | **Accession ID** |
| --- | --- | --- |
| 1. | CMV promoter + RBS | BBa_I712004 |
| 2. | LacR | BBa_K731500 |
| 3. | GFP+ Terminator | BBa_K259006 |
| 4. | Spacer DNA | BBa_K1123011 |
| 5. | pcDNA 3.1 (Backbone) | BBa_K3030004 |
